# The circulating immune cell landscape stratifies metastatic burden in breast cancer patients

**DOI:** 10.1101/2023.11.01.565223

**Authors:** S Mangiola, R Brown, J Berthelet, S Guleria, C Liyanage, S Ostrouska, J Wilcox, M Merdas, PF Larsen, C Bell, J Schroder, L Mielke, J Mariadason, S Chang-Hao Tsao, Y Chen, VK Yadav, RL Anderson, S Vodala, D Merino, A Behren, B Yeo, AT Papenfuss, B Pal

## Abstract

Advanced breast cancers show varying degrees of metastasis; however, reliable biomarkers of metastatic disease progression remain unknown. In circulation, immune cells are the first line of defence against tumour cells. Herein, using >109,591 peripheral blood mononuclear cells from healthy individuals and breast cancer patients, we tested whether molecular traits of the circulating immune cells, probed with single-cell transcriptomics, can be used to segregate metastatic profiles. Our analyses revealed significant compositional and transcriptional differences in PBMCs of patients with restricted or high metastatic burden versus healthy subjects. The abundance of T cell and monocyte subtypes segregated cancer patients from healthy individuals, while memory and unconventional T cells were enriched in low metastatic burden disease. The cell communication axes were also found to be tightly associated with the extent of metastatic burden. Additionally, we identified a PBMC-derived metastatic gene signature capable of discerning metastatic condition from a healthy state. Our study provides unique molecular insights into the peripheral immune system operating in metastatic breast cancer, revealing potential new biomarkers of the extent of the metastatic state. Tracking such immune traits associated with metastatic spread could complement existing diagnostic tools.

## Introduction

Breast cancer metastasises beyond the regional lymph nodes to distant sites, with variable locations and volumes of organ involvement [1]. Metastatic breast cancer (MBC) is generally incurable, with roughly 10-20% of patients developing fatal metastatic lesions at multiple sites, including lung, liver, bone and brain [2], [3]. One of the unresolved issues in the treatment and care of breast cancer patients is the timely detection of widespread distant metastasis. Compared with existing approaches (biopsies, imaging, etc.), blood-based cancer detection methods increase sample (tissue) accessibility and minimise invasive procedures. In recent years, efforts have been made to study blood-based biomarkers for cancer detection through qualitative and quantitative analyses of circulating tumour cells (CTCs) and circulating tumour DNA (ctDNA) [4–6].

Peripheral blood mononuclear cells (PBMCs), a component of the circulating immune cells, are considered the first line of defence against cancer [7] [8]. These cells are vital in responding to and being recruited to primary and metastatic tumour sites. Recent studies have shown that PBMCs contain cancer-specific biomarkers for the detection and prognosis of some cancer types, such as advanced renal cell carcinoma, non-small cell lung cancer and hepatocellular carcinoma [9–12]. Thus, exploring the diverse cellular composition and transcriptomic changes of PBMCs associated with metastasis could establish a molecular foundation and predictive biomarkers for the metastatic progression in cancer patients. In breast cancer, PBMCs of triple-negative breast cancer (TNBC) patients show compositional changes in immunosuppressive cell types and enrichment of a predictive gene signature (CD163, CXCR4, and THBS1) for relapse-free survival [13]. PBMC transcriptome profiles have also been used to subclassify breast cancer patients based on lymphocyte or neutrophil-polarised immune responses [14]. Whether these compositional and transcriptomic changes in PBMCs can be exploited as novel biomarkers of metastatic disease or hard-to-biopsy metastatic sites in breast cancer remains to be investigated.

Herein, we used single-cell RNA sequencing to investigate PBMCs isolated from ten patients diagnosed with metastatic breast cancer, with five classified as low metastatic burden patients and the other five with more extensive metastatic burden. By analysing the immune profiles of these patients, we aimed to establish a reliable molecular biomarker for identifying patients with low-volume metastasis amendable to localised therapies. Our study revealed a strong relationship between the single-cell transcriptomes of circulating immune cells and the extent of metastatic spread of breast cancer, thereby revealing unique biomarkers of metastatic burden. Our findings will assist in designing improved detection and treatment strategies against aggressive breast cancer and have significant implications for personalised cancer care and treatment decision-making.

## Results

### Characterising cellular heterogeneity in PBMCs of breast cancer patients

Blood samples of 10 female breast cancer patients (**Supp. Table 1**) at the time of diagnosis of their metastatic disease were used to isolate PBMCs. We used the droplet-based 10X Chromium workflow to capture single cells and prepare single-cell RNA sequencing libraries for PBMCs. The sequencing data was generated from 60,501 cells with a median of 3,776 per sample. Across samples, cells had a median RNA read output from 629 to 4,110 **(Supp. figure S1A)**. The rank of cells based on total RNA output (barcodeRanks; [15]) followed a nominal curve for all samples **(Supp. figure S1A)**. We excluded a median of 3% of all cells (across samples) from analysis due to high mitochondrial transcripts and a median of 6.7% of all remaining cells across samples as doublets.

Data integration and clustering analysis defined four major cell clusters **(Figure 1A),** corresponding to B cells, myeloid, plasmacytoid dendritic cells (pDC) and a lymphoid population of T cells and NK cells **(Figure 1B)**. Cell identity was inferred by mapping against a reference PBMC dataset [16] and with manual curation against known lineage markers **(Figures 1C and D)**. The T/NK cell (#4) and myeloid cell clusters (#2) were independently re-clustered and reanalysed (**Figures 1E-J)**. For the T/NK cluster, sixteen transcriptional subclusters were identified in the lymphoid cluster, including CD4+ T cells (six clusters), CD8+ T cells (five clusters), NK cells (three clusters), gamma delta (γδ) T cells, and <u>M</u>ucosal-<u>a</u>ssociated invariant <u>T</u> (MAIT) cells **(Figure 1E)**. All cells transcribed varying levels of CD3G, a pan T-cell gene, whereas CD4 cells co-transcribed IL7R and KLRB1, while CD8 cells were marked by GNLY and GZMK **(Figure 1E and F)**. The MAIT, NK and γδ T cells exclusively transcribed lineage markers NCR3, STMN1 and TRDC, respectively **(Figure 1F and G)**.

**Figure 1.**
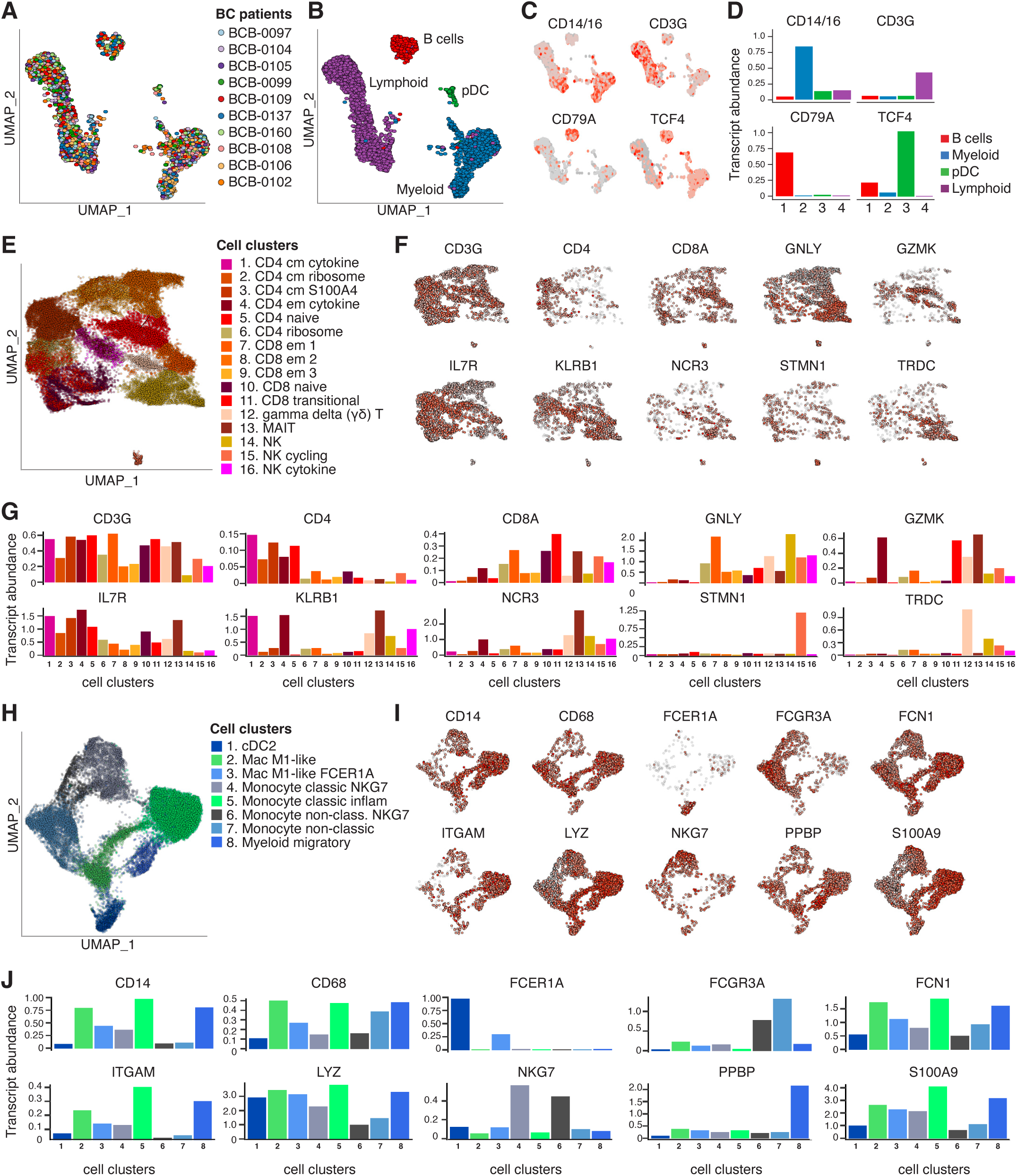
Mapping cellular heterogeneity in myeloid and lymphoid cell compartments of PBMCs in metastatic breast cancer patients. **A.** UMAP plot coloured by BC patient sample. **B.** UMAP plot coloured by four major cell types (B, T+NK, monocyte-derived, and plasmacytoid dendritic cells). **C.** UMAP plot coloured by the relative transcript abundance of marker genes for the four major cell types. **D.** Summary of the scaled relative transcript abundance of the marker genes for the four major cell types. **E.** UMAP plot of the lymphocytes re-analysed in isolation, coloured by cell type. **F.** UMAP plots coloured by the relative transcript abundance of marker genes for the lymphocyte cell types. **G.** Summary of the scaled relative transcript abundance of the lymphocyte marker genes. **H.** UMAP plot of the myeloid cells re-analysed in isolation, coloured by cell type. **I.** UMAP plots coloured by the relative transcript abundance of marker genes for the myeloid cells. **J.** Summary of the scaled relative transcript abundance of the marker genes for the myeloid cells.

The in-depth investigation of the myeloid population (**Figure 1B**) revealed eight cell sub-clusters consisting of cDC2, monocyte (four clusters), myeloid migratory cells and two unexpected clusters of macrophage-like cells **(Figure 1H)**. The cDC2 cells transcribe the high-affinity IgE receptor, FCER1A, while migratory myeloid cells co-transcribe high levels of chemokine PPBP **(Figure 1I and J)**. The monocytes and myeloid migratory cells showed varying transcription of CD14, ITGAM and CD68 lineage markers. While the classical and non-classical monocytes could be distinguished with differential expression of CD14 and FCGR3A, both subsets surprisingly showed high abundance of inflammatory NK cell granule protein, NKG7 **(Figure 1I and J)**.

### Peripheral immune landscape segregates healthy and cancer conditions

To identify molecular changes in the immune landscape that are associated with metastasis, we began by integrating publicly available PBMC single-cell RNA sequencing datasets of 10 healthy donors [17] [18] [19] [20] (**Supp Table 1**) with single-cell transcriptomes of patients with breast cancer (**Figure 2A, top panel)**. The healthy PBMC data represent 3 female and 7 male donors. The abundance of sex markers (XIST for females, DDX3Y for males) was confirmed (**Supp. figure S1C**). After filtering for empty droplets and dead cells, we selected 49,090 cells with a median of 4,137 per subject. The median RNA read output **(Supp. figure S1B)** for each healthy sample was comparable to cancer samples, with roughly 5,000 cells having at least 10,000 reads. The principal component analysis of the healthy and cancer cohort, using pseudobulk representation of the single-cell data, separated samples based on clinical characteristics (healthy vs. cancer) (**Figure 2A, bottom panel**). However, gender did not associate with the two principal components, indicating that using a mixed-gender healthy baseline will not be detrimental for subsequent analysis (**Supp. figure S1D)**.

**Figure 2.**
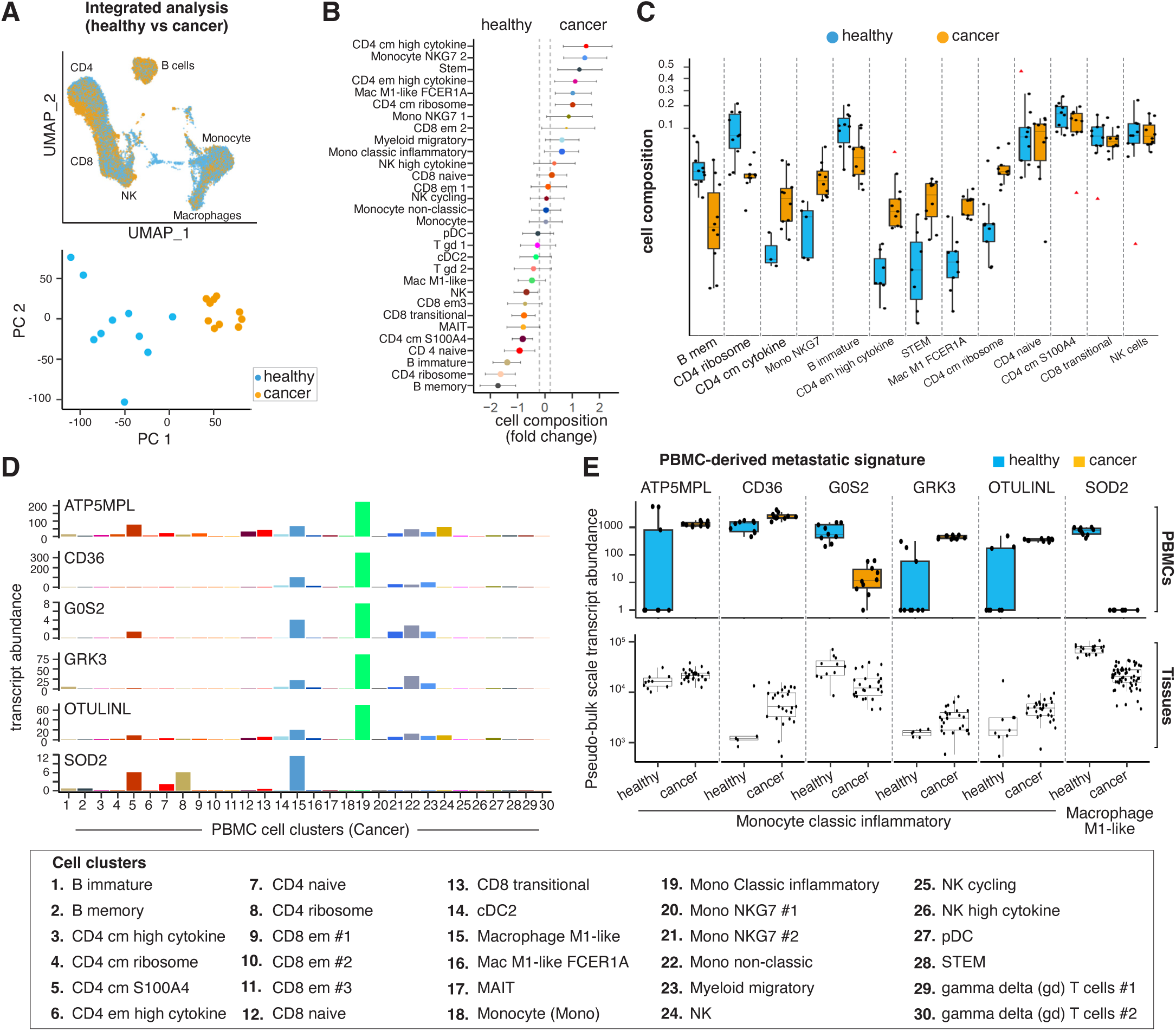
Metastatic breast cancer is associated with unique transcriptomic changes in circulating immune cells. **A-upper panel.** The integrated UMAP plot of ten healthy and ten cancer PBMC transcriptome profiles. The colour gradient indicates UMAP areas enriched for healthy (blue) or cancer (yellow). **A-lower panel.** PCA plot of the 20 pseudo-bulk transcriptomes, coloured by healthy (blue) and metastatic (yellow) conditions. **B.** Differential cell composition between healthy and cancer patients. Dots are coloured by cell type (see Figure 1, panels E and H). The right side of the plot contains cells that are enriched in cancer patients, while the cells on the left are enriched in healthy donors. The error bars represent the 95% credible interval for the estimates. The significantly enriched cell types are defined by having the 95% credible interval outside the −0.2/0.2 logit-fold-change interval. **C.** Cell type proportionality for the cell types ranked by their differential abundance. The red triangles represent outliers identified by sccomp. **D.** Transcript abundance per cell type (in cancer) for the six-gene signature differentiating healthy from cancer. **E upper panel.** Healthy/cancer gene signature. Boxplots represent the scaled transcript abundance distribution of the cell-type/sample-specific pseudo-bulk data, healthy (blue) and cancer-associated (orange). **E lower panel.** Validation of the PBMC signature in breast tissue immune cells.

The PBMC composition analysis revealed a monocytic- and T-memory-based inflammatory response in cancer, with a relative depletion of B cells (memory and immature) and CD4+ T cells (naïve, ribosome high) in cancer samples compared to the healthy cohorts (**Figure 2B and 2C**). On the contrary, CD 4+ T cells with high cytokine and central memory phenotypes were enriched in cancer samples (**Figure 2B and 2C**). Unconventional T cell populations, including MAIT and γδ T cells, which are known to kill tumour cells directly via MHC-independent mechanisms [21] [22] [23], were found to be present at decreased frequency in PBMC samples of metastatic breast cancer patients. In the myeloid compartment, the inflammatory monocytes (NKG7 high transcription), M1 macrophage-like cells (FCER1A high transcription), and myeloid cells with migratory phenotypes were enriched in metastatic breast cancer patients. Collectively, these results suggest that the blood of healthy donors and cancer patients contain distinct immune repertoires. These systemic changes in immune cell composition in cancer patients could potentially play a role in evasion and cancer progression.

### PBMC-derived transcriptomic signature correlates with breast cancer

We next tested whether changes in systemic immune profiles could provide translational opportunities. To this end, we sought to generate a PBMC-derived transcriptional signature that would segregate healthy individuals from cancer patients. We first identified the differentially expressed gene transcripts in all the 30 cell types of PBMCs shown in Figure 2A (**Supp. file 1**). Next, significantly differentially expressed genes (FDR < 0.05) were interrogated in the corresponding cell types of the single-cell transcriptomic profiles generated from 13 healthy breast tissues and 35 primary breast tumours representing major breast cancer subtypes, including estrogen receptor (ER)+, HER2+, and TNBC [24]. This analysis identified six genes, ATP5MPL, SOD2, CD36, G0S2, GRK3 and OTULINL, hereinafter referred to as PBMC-derived metastatic signature, that were differentially expressed between healthy and cancer conditions in both the PBMCs and the tissue microenvironment. The signature genes are expressed mainly by macrophage M1-like cells and inflammatory monocytic populations in PBMCs (**Figure 2D**). Interestingly, the upregulated (ATP5ML, CD36, GRK3, OTULINL) and downregulated (G0S2, SOD2) genes show similar differential expression patterns between healthy and cancer stages in the same immune cell types in both systemic (PBMCs; **Figure 2E, top panel**) as well as tissue-resident microenvironment (healthy breast vs tumour; **Figure 2E, bottom panel**).

### Metastatic progression is associated with the downregulation of anti-cancer immune surveillance and cytotoxicity

Next, we investigated the ability of the cellular and molecular characteristics of the peripheral immune system to resolve low or high-burden metastatic profiles by dichotomising our sample cohort for low (**single organ; blue**) or high (**multiple organs; red**) metastatic burden **(Supp. table 1 and Figure 3A)**. The PBMC composition analysis revealed differences associated with metastatic burden **(Figure 3B)**. The T cell compartment for CD4+ and CD8+ was significantly depleted in high-burden metastasis, including effector memory (em), central memory (cm), and transitional and high cytokine-producing cells. The relative depletion of unconventional T cells, including MAIT and gamma delta (γδ) T cells (Vδ 2 subtype), were among the strongest indicators of high-burden disease. In contrast, low-burden oligometastatic patients showed a systemic immune response skewed toward B cells (immature and memory). In the myeloid compartment, pre-dendritic, inflammatory classic monocytes and M1 macrophage-like cells were depleted in the high-burden metastasis group. Collectively, these immune cell populations are key drivers of the anti-tumour immune response.

**Figure 3.**
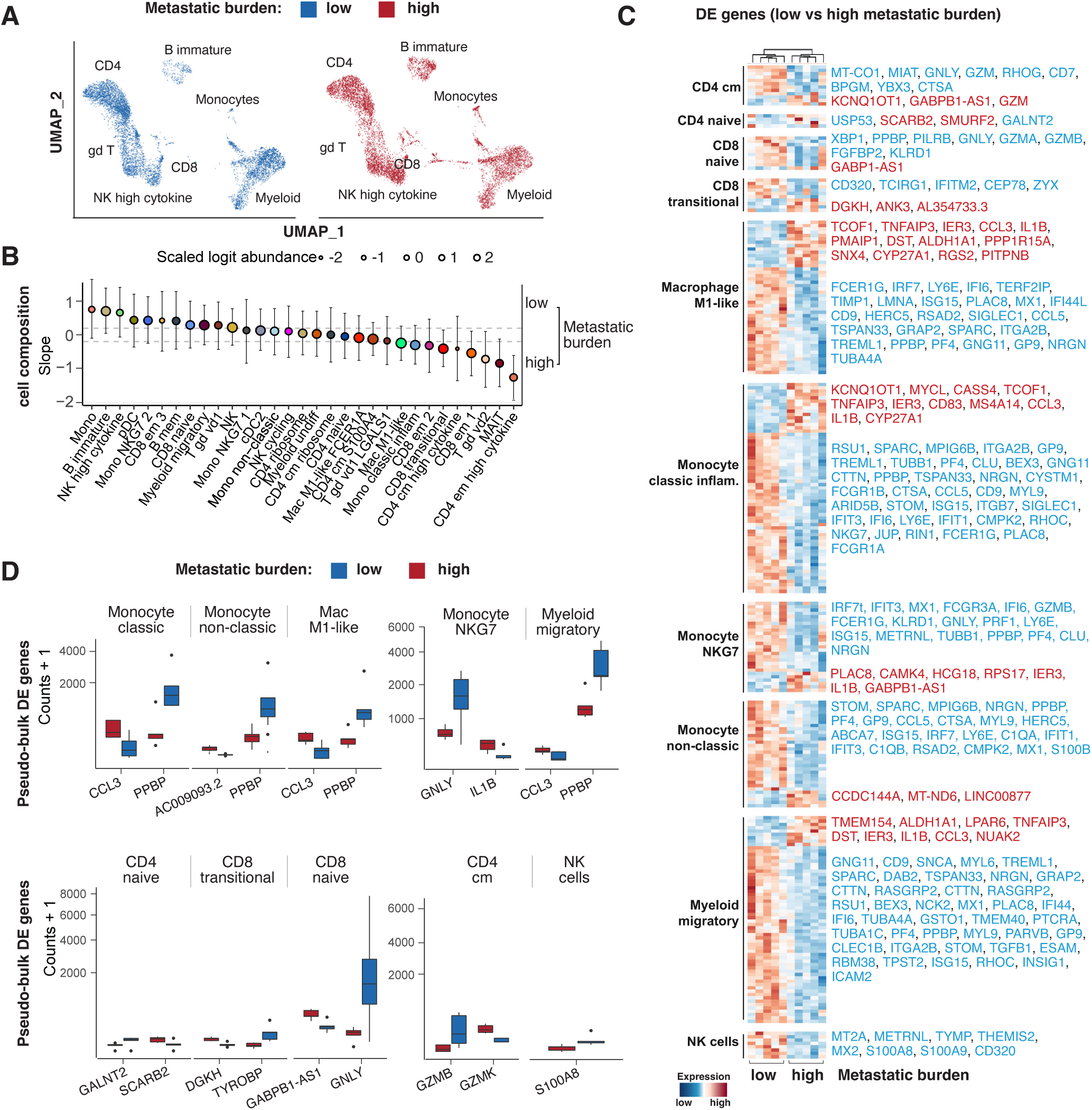
Metastatic burden redefines the cell composition and molecular characteristics of PBMCs. **A.** Integrated UMAP plots showing single cells coloured by low (blue) and high (red) metastatic burden. **B.** Differential cell composition analysis between low and high metastatic burden. Dots are coloured by cell type (see Figure 1, panels E and H). Cell types enriched in low-metastatic-burden patients are at the top, while those enriched in high-metastatic-burden patients are at the bottom. The error bars represent the 95% credible interval for the estimates. The bold dot borders label the significantly enriched cell types (testing for an effect bigger than 0.2 logit fold change). **C.** Scaled transcription abundance of cell-type-specific pseudo-bulk samples for the differentially abundant gene transcripts between low vs high metastasis burden. **D.** Boxplots showing the scaled transcript abundance distribution of the cell-type/sample-specific pseudo-bulk data for low (blue) or high (red) metastatic burden.

The enriched and depleted cell types also showed different transcriptional phenotypes between metastatic profiles (**Figure 3C**). The most significant transcriptional differences were observed in CD4+ T (central memory and naïve), CD8+ T (transitional and naïve), monocytes (classic, non-classic and NKG7 high), and migratory myeloid and macrophage M1-like cells (**Figure 3C**). In high metastatic disease, the lymphoid populations CD4+ and CD8+ T cells show upregulation of GZMK (pro-inflammatory) [25] and downregulation of GZMB and GNLY (cytolytic activity) [26] [27]. However, NK cells in high metastatic disease are marked by relatively lower expression of S100A8, a pro-inflammatory factor that promotes metastasis [28, 29]. In contrast, the myeloid cell types in high metastatic burden are marked by upregulation of CCL3 (acute inflammation) [30] and IL1B (acute inflammatory response) [31] [32] and downregulation of PPBP (glucose metabolism and plasminogen activator) [33] and GNLY (cytolytic) [27] (**Figure 3D**).

### Metastatic progression is associated with the disruption of key cell-cell communications in circulating immune cells

In the tissue microenvironment, intercellular crosstalk has been rigorously investigated [34] [35] [36]. Yet, the precise modalities governing communication among circulating immune cells in oncological contexts remain mostly unknown. To address this, we used CellChat [37] to investigate the co-transcriptional dynamics of ligand-receptor pairs across diverse cellular populations. Such exploration provides insights into both short (involving direct cell-to-cell contacts and matrix interactions) and long-range (mediated by secreted ligands) communication axes associated with metastatic burden.

A pronounced decrease in intercellular communication, including short- and long-range signalling, was observed in high metastatic disease compared to low metastatic burden **(Figure 4A)**. Conventional dendritic cells (cDC2) showed the most significant downregulation among all cell populations **(Figure 4B)**. The GALECTIN communication pathway was downregulated between cDC2 cells and the largest number of other cell types. This pathway includes LGALS9 as a ligand from dendritic cells and PTPRC, HAVCR2 and CD44 as receptors from target cell types **(Figure 4C)**. The Visfatin/NAMPT communication pathway was downregulated in fewer cell types, including inflammatory monocytes and dendritic cells. We also identified the downregulation of the secretory molecule Thrombospondin-1 (encoded by THBS) and its cognate receptors CD47 and CD36 [38] between monocytes and pDC2 for high metastatic burden (**Figure 4D**).

**Figure 4.**
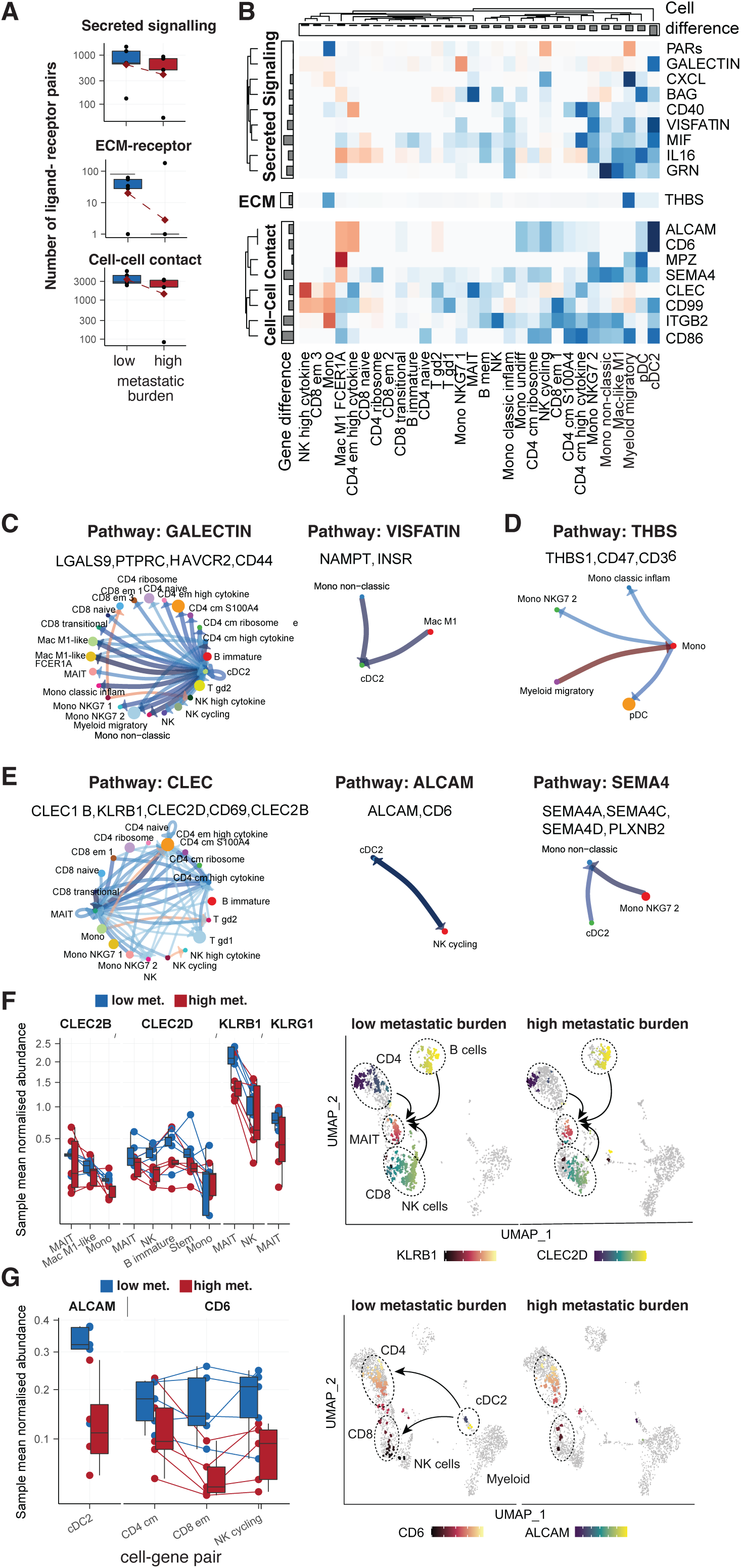
The extent of metastatic burden alters immune crosstalk profiles. **A.** Comparison of the number of highly transcribed ligand-receptor pairs between PBMC samples from high and low metastatic burden patients. The communication axes are grouped by long-distance secreted signalling, extracellular matrix (ECM) receptors, and cell-cell interactions. The red diamond indicates the average count across samples. **B.** Composite enrichment scores for cell-type/communication axes between a cell type (e.g. cDC2; x-axis) and all other cell types. Each communication axis can underlie several ligands and receptors. The colours represent the enrichment direction (blue for low and red for high metastatic burden disease). Communication axes are grouped by type. Bar plots represent the enrichment score average for cell types (columns) or communication axes (rows). **C, D, and E.** Enrichment of communication among cell type pairs for the most differentially enriched communication axes (in either direction). The colours represent the enrichment direction (blue for low and red for high metastatic burden disease). The genes included in each communication axis are below the names of the communication axes. The arrows represent the direction of the communication (ligand outgoing, receptor incoming). Some communications are bi-directional. **F.** Transcript abundance for the CLEC/KLRB1 ligand/receptor pair for the cell types that show enrichment. Colours represent conditions (blue for low and red for high metastatic burden). UMAP plot of cells coloured by transcription of either CLEC or KLRB1. The UMAP panels on the right side show the enrichment of communication between cell types for the most differentially enriched communication axes (in either direction). **G.** Transcript abundance for the ALCAM/CD6 ligand/receptor pair for the cell types that show enrichment (dendritic cells with ligand and lymphocytes with the receptor). Colours represent conditions. UMAP plot of cells coloured by transcription of either ALCAM or CD6. The UMAP panels (right-side) show communication enrichment among cell types. The colours represent the enrichment direction (blue for low- and red for high-metastatic burden disease).

For short-range interactions, the communication pathway ALCAM and CD6 showed the strongest downregulation in high-burden metastasis **(Figure 4E and G)**. Mucosal-associated invariant T cells (MAIT) showed several depleted communication axes, such as CD69, CLEC, and KLRB1 (Figure 4E and F). Within this network, high cytokine-expressing CD4 memory and γδ T cells emerge as key interactive hubs. Amongst myeloid cells, the SEM4A communication pathway, known to play a role in the T-cell priming [39], was significantly downregulated for high metastatic burden **(Figure 4E)**.

### Circulating unconventional T cell molecular profile changes with metastatic burden

The pathogenic roles of MAIT and γδ T cells have been studied extensively [40] [41]. However, their role in breast cancer and metastasis is unclear. Interestingly, our PBMC analysis revealed depleted numbers of MAIT and γδ T cells in samples associated with high metastatic burden compared to healthy controls or low metastatic burden conditions (**Figures 2B and 3B**). Hence, we re-clustered these cells from total PBMCs for further in-depth analysis.

MAIT cells form a relatively homogeneous cell cluster in healthy and cancer conditions (**Supp. figure S2A and B**). In cancer patients, the MAIT cells show upregulation of NCR3 (cytotoxicity triggering receptor) [42], ARL4C (ADP-ribosylation factor-like protein associated with cancer progression) [43], LYAR (a negative regulator of the innate immune response) [44] [45] and DUSP2 (a MAP kinase family member required for cellular differentiation and proliferation) [46] (**Supp. figure S2C**). Whereas the healthy MAIT cells are marked by relatively higher expression of genes, including LTB, known to regulate proinflammatory response and development of lymphoid tissue [47] and, S100A4 involved in motility, recruitment and chemotaxis of inflammatory immune cells [48] (**Supp. figure S2C**).

Compared to MAIT cells, the γδ T cells show distinct transcriptional heterogeneity in healthy and cancer conditions (**Supp. figure S3A and B)**. We interrogated re-clustered γδ T cells with previously described 30-gene signature [49] to identify Vδ1 and Vδ2 subsets in γδ T cell population of healthy and cancer PBMCs (**Supp. figure S3A and B, left panels and S3C)**. As expected, Vδ1 and Vδ2 subsets express genes involved in T-cell mediated cytotoxicity (GNLY, GZMH, FGFBP2, GZMB, NKG7, LGALS1) and proinflammatory responses (GZMK, LTB, IL7R). Next, we determined the differentially expressed genes that revealed unique molecular differences within γδ T subsets **(Supp. figure S3A and B, right panels)**. Especially in cancer samples, Vδ1 subset express SH3BGRL3, CYBA, CCL5, CST7, TMSB10, FCGR3A, ZEB2, PRF1, IFITM2, HLA-E genes, whereas Vδ2 cells are marked by CD52, TPT1 and EEF1A1 genes.

To further delineate the γδ T cell phenotypic characteristics associated with metastasis, we interrogated the Vδ1 and Vδ2 subsets in the context of low and high metastatic cases (**Figure 5A**). The Vδ1 cell cluster (2; yellow) was relatively enriched in PBMCs from patients with high metastatic burden compared to the Vδ2 cell cluster (3; green) (**Figure 5B**). The Vδ2 cell cluster showed minor changes in the expression of proinflammatory genes IL7R, GZMK and LTB between low and high metastatic conditions (**Figure 5C**). Whereas the Vδ1 subsets expressing cytolytic genes (FGFBP2, GNLY, GZMB, GZMH, LGALS1 and NKG7) were detected at increased frequency in patients with high metastatic burden **(Figure 5C)**. Therefore, we speculate that high-burden multi-organ metastasis drives phenotypic and compositional changes in circulating γδ T cell subsets. Whether metastasis orchestrates similar phenotypic changes in tissue-resident γδ T cells remains to be determined. To this end, we completed multiplex immunohistochemistry (mIHC) on primary tumour tissues from 6 non-metastatic early breast cancer (EBC) and 4 metastatic breast cancer (MBC) patients to determine the relative proportion of FGFBP2^+^ γδ T cells between these two conditions. The primary tumour tissues did not show significant changes in total γδ T cells numbers; however, the tissue-resident FGFBP2+ γδ T cells were significantly enriched in MBC tumours (**Figure 5D and E**), suggesting that molecular mechanisms similar to the peripheral immune system may operate in primary tumour microenvironment (TME) to regulate the recruitment of cytotoxic γδ T cell subsets.

**Figure 5.**
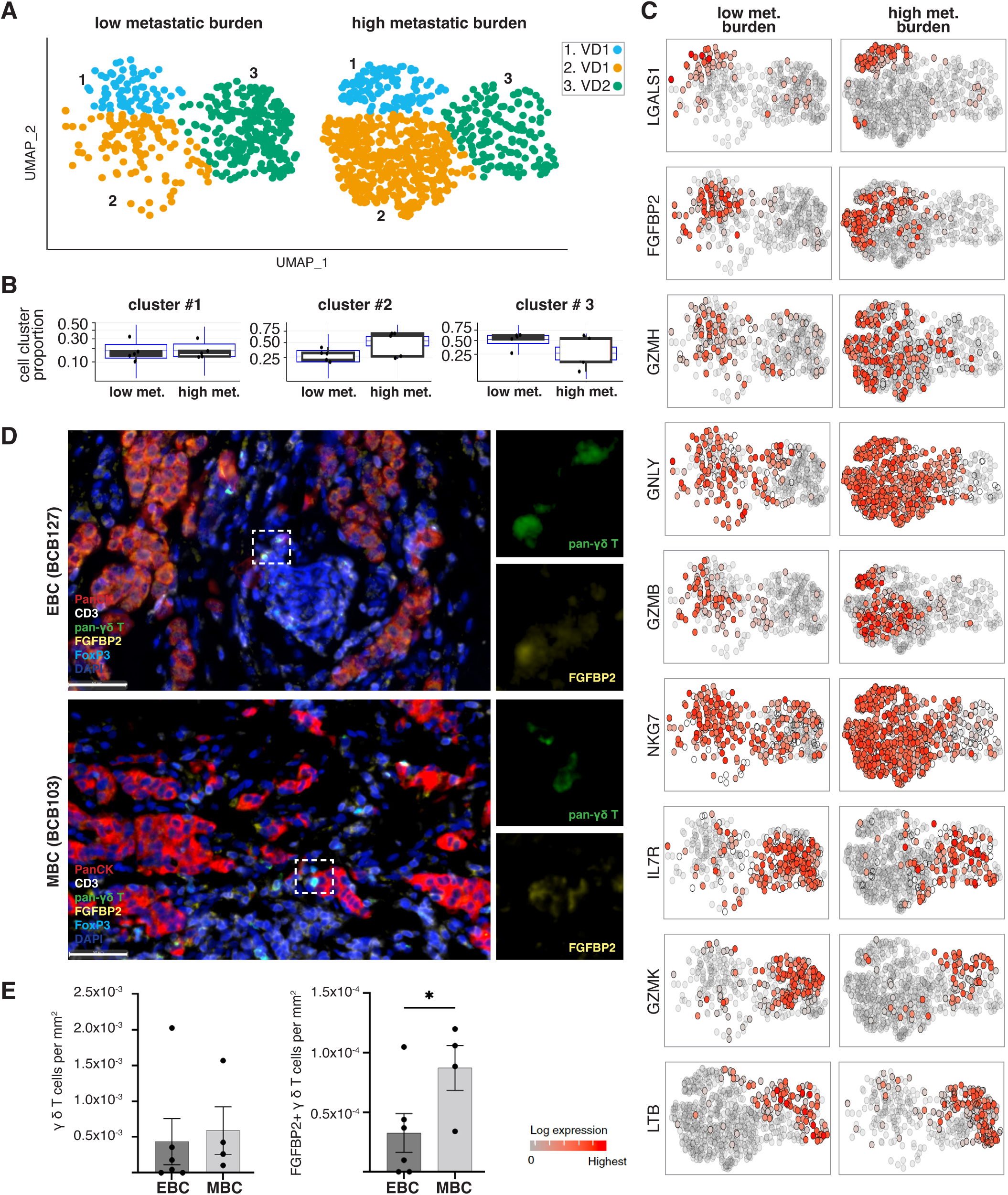
Gamma-delta T cells show transcriptional changes in metastatic patients with breast cancer. **A.** UMAP plots of the re-clustered γδ T cells from low-metastatic (left panel) and high-metastatic burden samples (right panel). **B**. Differential composition (estimated by sccomp) of the three gamma-delta cell clusters relative to the whole gamma-delta population between low and high-burden metastatic patients. Data points are samples. The blue boxplot represents the posterior predictive estimates that indicate the model’s descriptive accuracy for the data [82]. **C**. UMAP plots of gamma-delta T cells in low-metastatic (left panel) and high-metastatic samples (right panel), coloured by the relative transcript abundance of the top 9 marker genes. **D.** mIHC panel shows the expression of pan-cytokeratin (red; tumour cells), CD3 (white; T cells), pan-γδ T cell (green), FGFBP2 (yellow), FOXP3 (light blue) and DAPI (dark blue; nuclei) in primary tumour tissues from non-metastatic early breast cancer patients (EBC; top panel) and metastatic breast cancer patients (MBC; bottom panel). The white dotted box mark γδ T cells that co-express FGFBP2 and are shown in magnified images on the right side. **(E)** mIHC image quantification showing the total number of γδ T cells per mm^2^ of tissues (left panel), and FGFBP2+ γδ T cells (right panel) from EBC (n=6) and MBC tumours (n=4) with p value (* = 0.0369).

## Discussion

Our study provides the first comprehensive analysis of single-cell transcriptomes of peripheral blood mononuclear cells (PBMCs) of breast cancer patients with metastatic disease. Data analysis revealed four major cell clusters corresponding to B cells, myeloid, plasmacytoid dendritic cells (pDC) and a lymphoid population comprising T and NK cells. Sixteen transcriptional subclusters were identified in the lymphoid cell compartment, including unconventional T cells and eight subclusters in the myeloid cell compartment. Interestingly, the classical and non-classical monocytes transcribe the inflammatory NK cell granule protein, NKG7. The NKG7 is required for NK and CD8^+^ T cell cytotoxic degranulation and CD4^+^ T cell activation and pro-inflammatory responses [50]. However, its functional relevance in monocyte activity needs further investigation.

By comparing single-cell transcriptome profiles from healthy individuals and patients with metastatic breast cancer, we identified novel PBMC-derived metastatic gene signatures potentially capable of detecting metastatic disease via blood analysis. The PBMC-derived metastasis signature comprises six marker genes, including ATP5MPL, CD36, GRK3, and OTULINL, that are upregulated in cancer, and SOD2 and G0S2 genes are highly expressed in healthy individuals. Overall, the PBMC-derived metastatic signature genes seem to have a critical role in cell proliferation and inflammation. The G0/G1 switch gene 2 (G0S2) has critical functions in cell survival and metastasis of cancer cells [51] [52] and in mediating a pro-inflammatory process by circulating monocytes [53]. ATP5MPL is required to maintain ATP synthase activity in the mitochondria [54]. ATP synthesis by mitochondria is critical for T cell proliferation, differentiation and effector function in cancer [55]. SOD2 encodes a mitochondrial protein required for radical scavenging to balance the intracellular levels of reactive oxygen species (ROS) [56]. CD36 is a transmembrane glycoprotein expressed by monocytes and acts as a scavenger receptor to mediate pro-tumour functions [57]. Among the top significant genes detected **(Figure 2E**), the human protein atlas predicts ATP5MPL as a prognostic ovarian cancer marker. However, its role during metastasis is not known. Polymorphisms in SOD2 are associated with an increased risk of non-Hodgkin lymphoma and lung and colorectal cancers [58]. GRK3 belongs to a subfamily of G protein-coupled receptor kinases that promote cancer progression by mediating cancer cell proliferation and is a poor prognostic indicator in multiple cancers [59] [60]. OTULINL is involved in the deubiquitination of poly-ubiquitin chains and plays an important role in autoimmunity, inflammation and infection [61] [62].

Overall, the metastatic gene signature mainly consists of inflammatory monocyte-specific markers robustly translated in circulating PBMCs and within the primary breast tumours of all major subtypes [24]. Whether these peripheral immunological changes are driven by or promote metastasis remains to be resolved. A previous study utilising mass spectrometry-based proteomics has shown phenotypic similarities between PBMCs and the TME of matching breast cancer tissues [63], whereby the expression profiles of IL-17, PI3K-Akt, and components of HIF-1 signalling pathways were found to be conserved between circulating and tumour infiltrating immune cells.

Further analysis comparing patients with metastatic disease revealed depletion of circulating CD4 and CD8 T effector memory cells, MAIT and γδ T cells in high metastatic burden cases. This pattern points toward possible unknown mechanisms operating in patients with low-burden metastatic disease that establish a more effective adaptive immune response and protection against the metastatic spread. The systemic immunity of low and high-burden disease is also characterised by distinct molecular changes in the lymphoid and myeloid populations, including monocytes, NK, and CD4+ and CD8+ T cells. Whether these phenotypic alterations in systemic immune cells and their compositional change are responsible for metastatic disease progression remains to be investigated.

Our data indicate that the circulating levels of unconventional T cells, including γδ T cells, are depleted during cancer and disease progression. However, in high metastatic burden disease, the circulating Vδ1 subset shows higher transcription of cytotoxicity genes NKG7, GZMH, GZMB, LGALS1 (Galectin 1) and FGFBP2 indicating a potential functional role of circulating γδ T cells during high burden metastatic disease. Interestingly, the total γδ T cell population is depleted in metastatic TME, pointing towards common mechanisms operating during metastasis that downregulate γδ T cells in the peripheral and tumour-local immune system. An integrated functional analysis of circulating and tumour resident γδ T subsets will be required to allow the design of efficient therapeutic strategies to boost γδ T cell activity against metastatic disease.

At the global transcriptome level, we observed downregulated cell-cell communication for high metastatic burden, especially between myeloid and lymphoid compartments. This included the short- (i.e., cell-cell contact, extracellular-matrix) and the long-range (secreted signalling) pathways. Key altered communication axes were ALCAM/CD6, between DC and T/NK cells, essential for dendritic/T cell effective interactions [64]; CD69, which is a hallmark of activated MAIT cells [65], which is also linked to the presence of CD25, the degranulation indicator CD107a, the generation of cytotoxic elements like perforin and granzyme B, and the emission of pro-inflammatory cytokines such as IFN-γ, TNF, IL-17, and CSF2/GM-CSF [65]; SEMA4/CD72, necessary for monocyte-promoted T cell activation [66]; and PECAM1 and ICAM, crucial to blood barrier migration [67]; and LCK, necessary for natural killer cell activation [68]. Additionally, key long-range immunomodulatory changes observed in lymphoid and myeloid compartments included the downregulation of cytokines (interferon-α/gamma and TNF-a pathways). Together, these findings uncover potential physiological functions of these phenotypic changes in circulating immune cells in metastatic disease and may provide a molecular basis of disease progression from low to high metastatic burden.

Our study focussed on metastatic burden and therefore further investigations into the role of tumour volume in influencing the immunological changes in the peripheral immune landscape are required. However, our data provide a strong rationale for using systemic immune cell composition and their marker genes to stratify metastatic breast cancer.

In addition to circulating tumour DNA (ctDNA), PBMCs offer unique advantages in accessibility, sensitivity and real-time assessment of tumour dynamics. Unlike imaging techniques that may have limitations in detecting microscopic lesions [69] [70] [71], systemic PBMC composition and marker genes can offer a minimally invasive approach to capture comprehensive molecular information about tumours. While ctDNA analyses provide a snapshot of the mutational landscape of tumour cells, PBMC profiling allows for an indirect evaluation of the local immune system at both primary and metastatic sites. Therefore, PBMC-based biomarkers will facilitate timely diagnosis of breast cancer or disease relapse, and treatment response. We speculate that the findings from our study can be leveraged to study other metastatic cancers. Furthermore, therapeutic targeting of cell-cell communication pathways in PBMCs may also lead to new immunotherapy treatments against metastatic disease.

## Material and Methods

### Human samples

Human breast tissues and blood samples were obtained from consenting patients through the Austin Health, a tertiary cancer hospital. Human Ethics approval was obtained from the Austin Human Research Ethics Committee. Clinicopathological characterisation, including site of metastasis are provided in **supplementary tables**.

### 10x Genomics Chromium library construction and sequencing

A 10x Genomics Chromium machine was used for single-cell capture and cDNA preparation according to the manufacturer’s Single Cell 3’ Protocol. The silane magnetic beads and Solid Phase Reversible Immobilization (SPRI) beads were used to clean up the GEM reaction mixture, and the barcoded cDNA was then amplified in a PCR step. The optimal cDNA amplicon size was achieved using the Covaris machine before library construction. The P7 and R2 primers were added during the GEM incubation, and the P5 and R1 during library construction via end repair, A-tailing, adaptor ligation and PCR. The final libraries contain the P5 and P7 primers used in Illumina bridge amplification. Sequencing was carried out on an Illumina Nextseq 500.

### Single-cell read mapping and quality control

The read information was converted to FASTQ files using bcl2fastq, mapped to the Hg38 3.0.0 genome and quantified using CellRanger [72] (version 7.0.0) with default parameters (--expect-cells=7000). The gene counts were imported to a SingleCellExperiment object [73] using DropletUtils [74] and manipulated using tidySingleCellExperiment (Bioconductor). Quality control and filtering were performed for each sample independently. The empty droplets were identified using barcodeRanks [74] below the inflection point. The mitochondrial gene names were downloaded using AnnotationDbi (mapIds) and EnsDb.Hsapiens.v86. Scuttle [75] was used for calculating cell quality metrics (perCellQCMetrics) with default parameters. The function isOutlier was used with default parameters to label outlier cells based on the fraction of mitochondrial transcription and ribosomal genes. For subsequent quality control and analyses, the data was converted to a Seurat container, and tidyseurat [76] was used for data manipulation. DoubletFinder (v3) [77]was used with default parameters to identify and filter out doublets.

### Bioinformatic analyses: normalisation, integration, and annotation

We identified variable genes within each sample using Seurat [78]. We estimated the cell cycle phase using CellCycleScoring. Using principal component analysis (PCA) and UMAP dimensionality reduction, we checked that the cell cycle was not a major confounder for cell distribution and grouping. We scaled counts cell-wise and removed the unwanted variation of residual mitochondrial and ribosomal content using the SCT transform [78]. The unwanted sample-wise technical variation was removed using the anchor-based integration implemented in Seurat [79], using SI-GA-E5 as the reference sample (as it was of excellent quality according to analyses and sequencing output). A nearest-neighbour graph is calculated from adjusted counts, and cells are clustered [80]. For the first-instance classification, we transferred cell-type annotation from the reference PBMC Azimuth dataset using Seurat [81].

We then isolated the macro clusters lymphoid, myeloid, and b-cells for independent integration, clustering, and marker gene selection. For each macro cluster, Seurat was used to identify cluster marker genes (FindAllMarkers, with the parameters only.pos = TRUE, min.pct = 0.25, thresh.use = 0.25). The curated cell annotation was based on a consensus between the identified markers and the label transfer described above (using pbmc_multimodal as the reference). The classification of the gamma delta population was refined using an ad hoc signature [49]. This signature included markers for gamma delta T cells vd1 and vd2. The gamma delta cells were further isolated, re-clustered (3 clusters found), and annotated with the ad hoc signature. Based on the cluster gene transcription abundance. The annotated macro clusters were then merged into a unique dataset. The healthy control dataset was taken from public resources GSE115189 [17], SCP345 (singlecell.broadinstitute.org), SCP424 [18], SCP591 [19], SRR11038995 [20], SRR7244582 [17], 10X 6K and 8K datasets from illumina.com, and annotated using our annotated dataset as the reference, as we have full control over the quality and content of our data. This choice was made to lower the risk of missing key cancer-related populations when integrating healthy individuals.

### Differential tissue composition

The method sccomp [82] was used for differential composition and variability analyses with the model ‘~ burden’. Predictive posterior intervals were checked after fit to assess the descriptive adequacy of the model to the data. Outlier observations (cell type abundance) were identified by sccomp and dropped from the hypothesis testing.

### Differential gene transcription abundance

The differential analyses were performed on pseudo-bulk sample/cell-type pairs, with the model ‘~ burden’. The framework tidybulk [83] was used to perform the analyses based on the robust edgeR implementation [84]. The hypothesis tested log 2-fold changes bigger than 1 [85]. The metastatic condition (non-metastatic, low or high-burden) was chosen as the only covariate. We detected outlier sample-gene observations using ppcseq [86]. The significant genes that included outliers were filtered out.

### Differential cell communication

The framework CellChat [37] was used for inferring the transcriptional abundance of ligand-receptor pairs between cell types. The transcriptional abundance of ligand-receptor pairs between cell types was estimated for each sample independently (with default parameters). The centrality of communication axes was estimated. We calculated the mean across samples of the same metastatic profile (low or high-burden) of the transcription score of ligand-receptor pairs between cell types. We then calculated the difference in the means to estimate the overall difference in communication. We confirmed the most significant communication changes at the gene count level. We created pseudobulk count aggregating cells at the sample/cell-type level. We selected the involved genes for each communication axis and tested the difference in pseudobulk transcription levels. Transcriptional levels were scaled for sample sequencing depth and sample-wise cell type abundance using TMM [87].

### Miscellaneous

We used the colour-deficiency-friendly code from dittoSeq [88]. For visualisation, we used mainly ggplot2 [89]. For heatmaps, we used tidyHeatmap [83] [90]. For data wrangling, we used the tidyverse software suite [91]. For pseudobulk differential transcript abundance analyses, we used tidybulk [83].

### Fluorescent multiplex-immunohistochemistry (mIHC)

Breast tumour sections were stained with antibodies using the Opal 7-7-colour multiplex immunohistochemistry kit (NEL811001KT, Akoya Biosciences). Staining was performed manually per the manufacturer’s protocol, as previously described (79). The following antibodies were used: CD3 (SP7, Invitrogen, 1:150 dilution), TCRδ (H-41, Santa Cruz Biotech, 1:100 dilution), pan Cytokeratin (AE1/AE3+5D3, Abcam, 1:200 dilution), FGFBP2 (polyclonal, Invitrogen, 1:700 dilution), FoxP3 (236A/E7, Abcam, 1:300 dilution) and spectral DAPI (Akoya Biosciences).

Whole slide scans at 10x magnification were performed using the Perkin Elmer Vectra Imaging System to mark regions of interest to obtain 20x magnification multispectral images (MSI) for analysis. A spectral library was created on InForm Analysis Software (Akoya Biosciences) for the quantitative analysis of cell populations. Immune cell populations of interest were identified by each marker’s mean fluorescence intensity (MFI), and the algorithm was applied to all samples. Images were analysed on HALO (Indica Labs) using Highplex v4.1.3 module for target cell phenotyping.

## Data availability

This study’s single-cell RNA sequencing data are available as the GEO series GSEXXXXX.

## Author contributions

SM, ATP, BY, and BP designed the study and analysed the data; SM, RB, JB, SG, CL, SO, JW, MM, CB and PFL performed the experiments; AB, JM, LM, SV, CY, YVK, STH, DM and RLA assisted in manuscript preparation.

---

**Supplementary table 1.** 10X single-cell library information.

| **Sample label** | **Metastatic burden/ site** | **Sum counts** | **Median counts** | **Median features** |
| --- | --- | --- | --- | --- |
| **BCB-0097** | Low - liver | 6977726 | 1326 | 705 |
| **BCB-0099** | Low - liver | 9477448 | 2566.5 | 1040.5 |
| **BCB-0109** | Low - liver | 20105016 | 3188 | 1155 |
| **BCB-0104** | Low - skin | 8703971 | 2826 | 1152 |
| **BCB-0105** | Low - mediastinal lymph node | 12594050 | 3637 | 1377 |
| **BCB-0137** | high - Axilla, SC LNs, soft tissues | 13428707 | 1342 | 688 |
| **BCB-0160** | high - clivia, femur, thoracic and lumbar spine, ribs | 11045516 | 1069 | 500 |
| **BCB-0108** | high - Multiple subcutaneous skin lesions, Pleura, Peritoneum/Omentum | 9686781 | 3299.5 | 1223.5 |
| **BCB-0106** | high - Axilla, lungs, Liver, Bone (multiple), Pleura, Skin | 16769560 | 629 | 364 |
| **BCB-0102** | high - Liver (multiple), Bone (mutiple, spine, femur, ribs) | 10150130 | 4109.5 | 1384 |
| **10x_6K** | healthy | 9884280 | 1926 | 730 |
| **10x_8K** | healthy | 35500342 | 4116.5 | 1305 |
| **GSE115189** | healthy | 6100426 | 2378 | 861 |
| **SCP345_580** | healthy | 21112154.52 | 3537.99 | 890 |
| **SCP345_860** | healthy | 24895466.59 | 3743.76 | 947 |
| **SCP424_pbmc1** | healthy | 14178849 | 4616.5 | 1516 |
| **SCP424_pbmc2** | healthy | 8759131 | 2663 | 1120 |
| **SCP591** | healthy | 1027974.21 | 1686.20 | 826 |
| **SRR11038995** | healthy | 35146551 | 3148 | 895 |
| **SRR7244582** | healthy | 10110503 | 3158 | 994 |

**Supplementary table 2.**
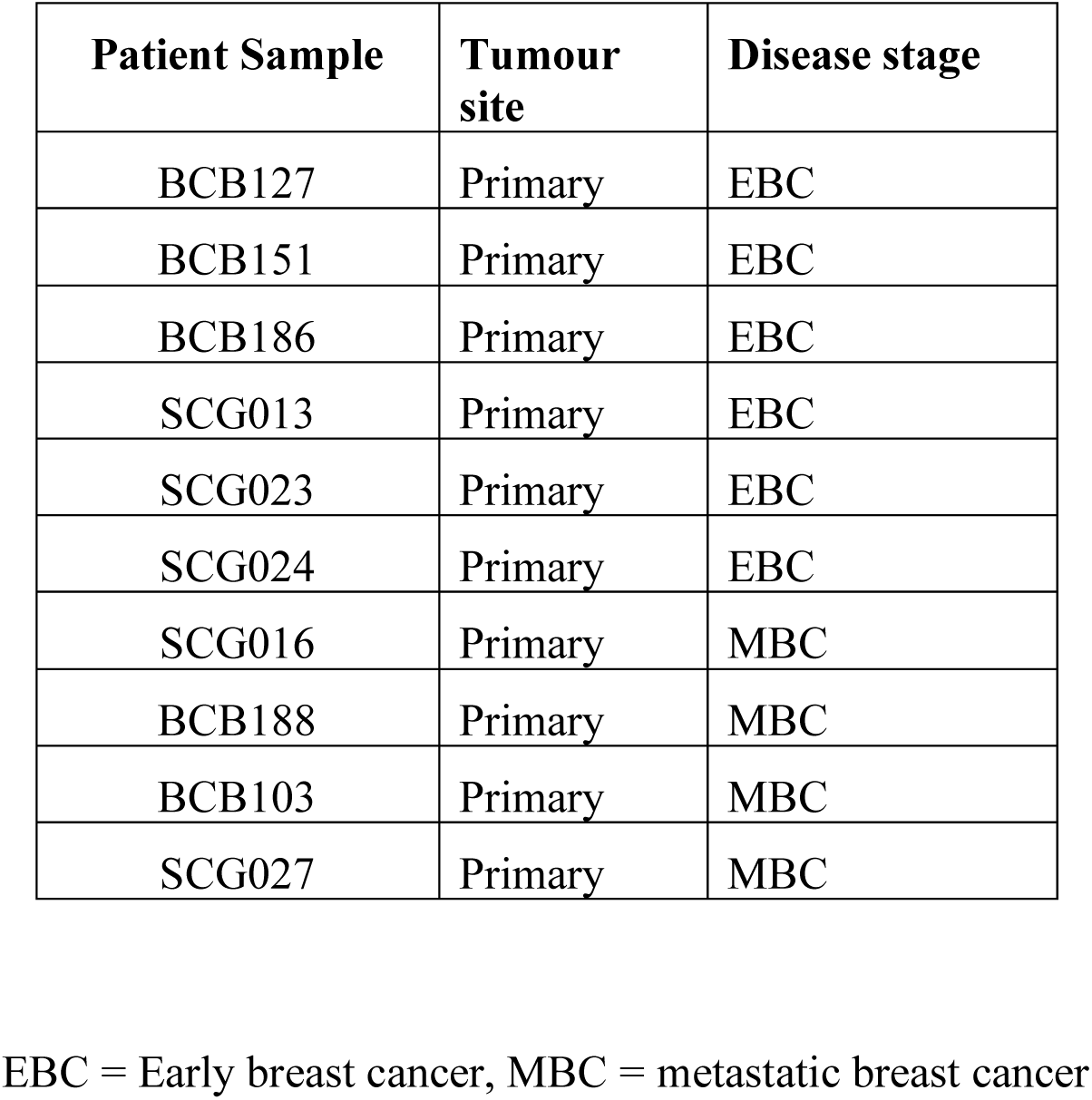
Breast tumour samples used for mIHC.

| **Patient Sample** | **Tumour site** | **Disease stage** |
| --- | --- | --- |
| BCB127 | Primary | EBC |
| BCB151 | Primary | EBC |
| BCB186 | Primary | EBC |
| SCG013 | Primary | EBC |
| SCG023 | Primary | EBC |
| SCG024 | Primary | EBC |
| SCG016 | Primary | MBC |
| BCB188 | Primary | MBC |
| BCB103 | Primary | MBC |
| SCG027 | Primary | MBC |

EBC = Early breast cancer, MBC = metastatic breast cancer

**Supplementary Figure S1.**
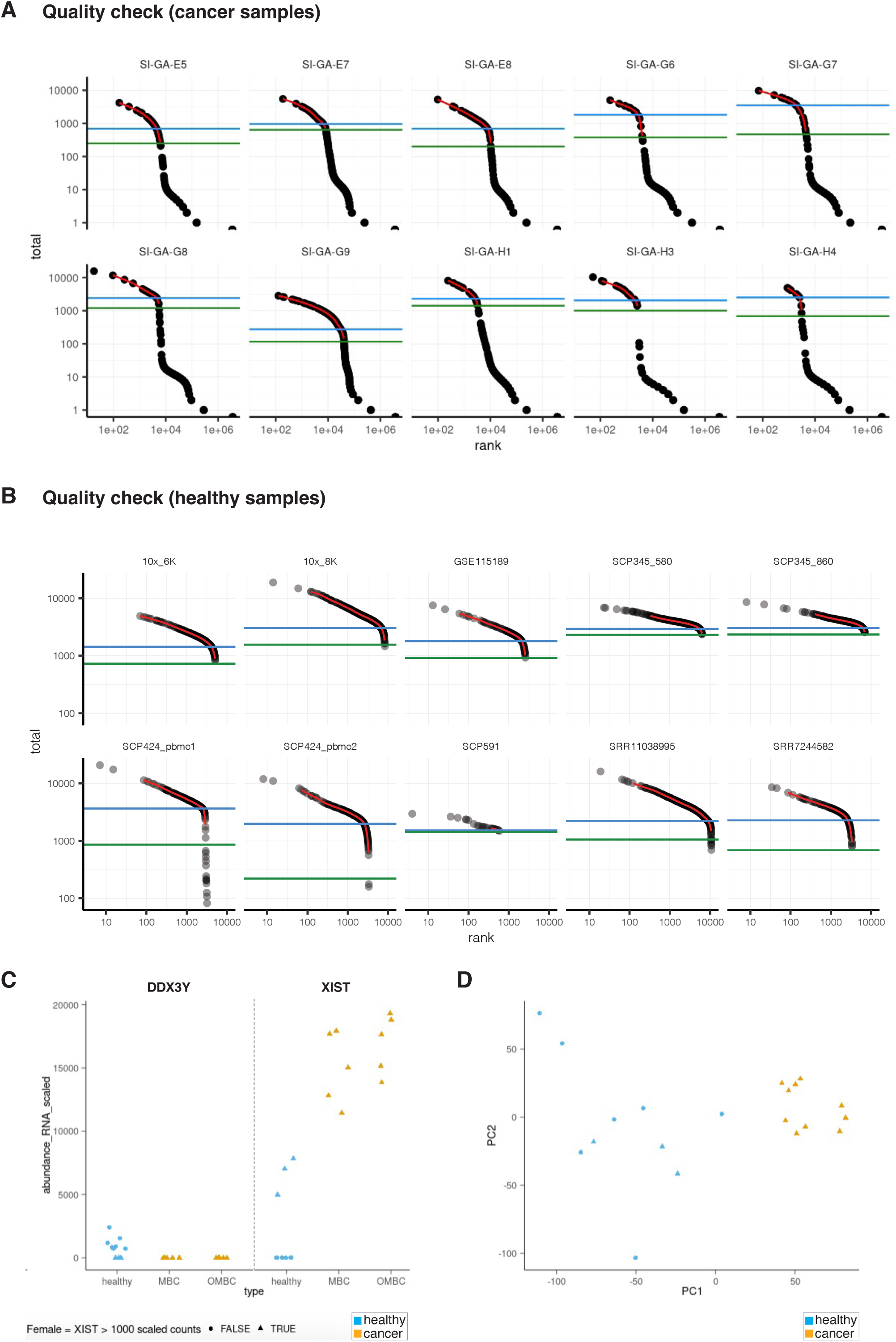
Quality check analysis of healthy and cancer single-cell transcriptome profiles. **A-B.**Ranks of cells by their total RNA counts. These plots were used to detect empty droplets according to the green inflection line. **B.** The use of a mixed-sex baseline does not have detrimental effects. The abundance for two sex markers (XIST for females, DDX3Y for males). The heuristic threshold of 1000 scaled read count for XIST classifies females from males. **C.** Principal component analysis of the healthy and cancer cohort, using pseudobulk representation of the single-cell RNA sequencing data. The first principal component separates clinical characteristics, while sex is not associated with the two principal components.

**Supplementary Figure S2.**
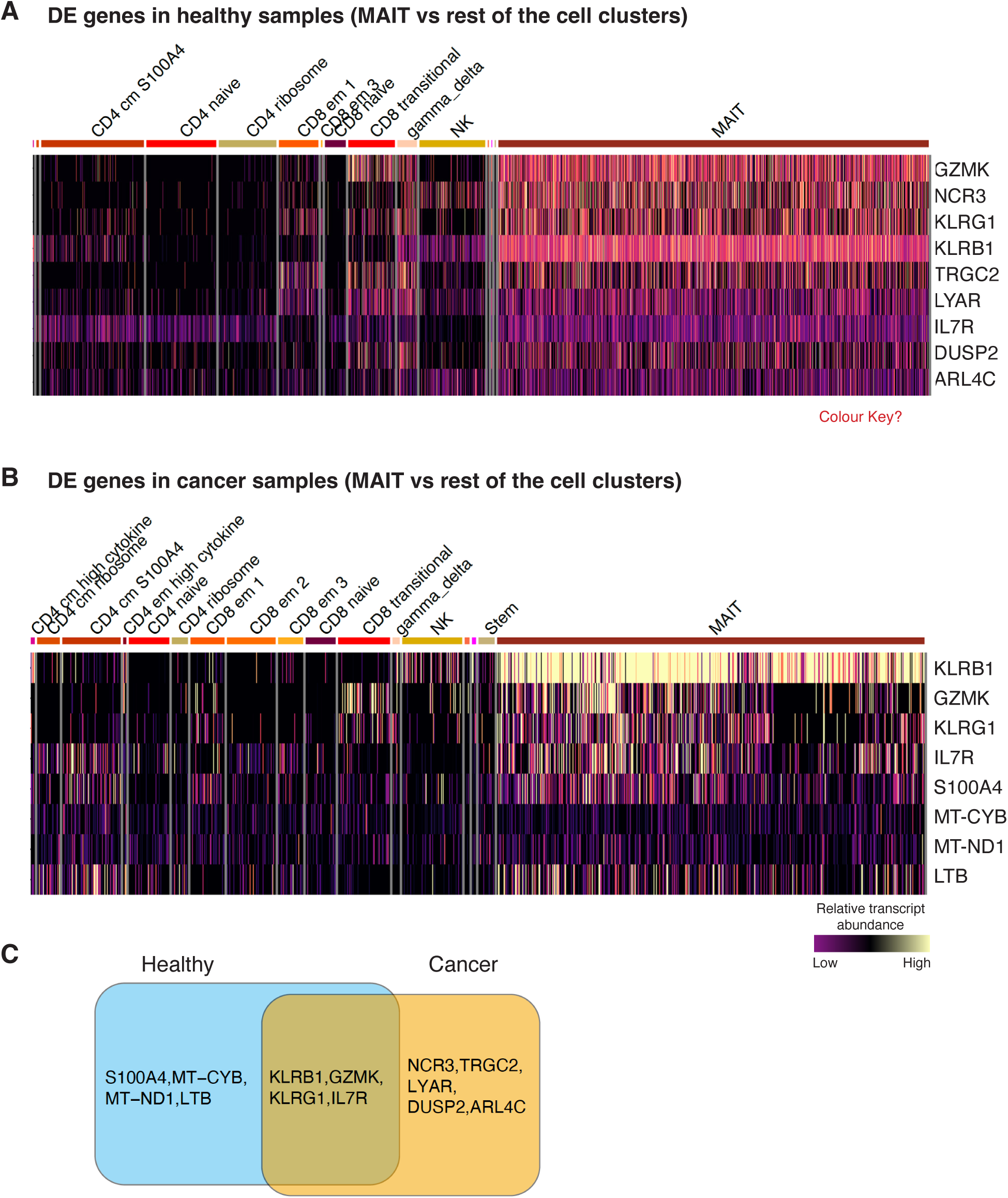
Gene expression analysis of circulating MAIT cells in healthy and breast cancer samples. **A-B.** Heat map showing top differentially abundant gene transcripts for MAIT cells compared to other lymphocytes in healthy and cancer samples. MAIT cells were re-clustered from Figure 2A. The heatmap is coloured by the row-scaled relative abundance of marker-gene transcripts. **C.** Top marker genes of MAIT cells in healthy and cancer cohorts.

**Supplementary Figure S3.**
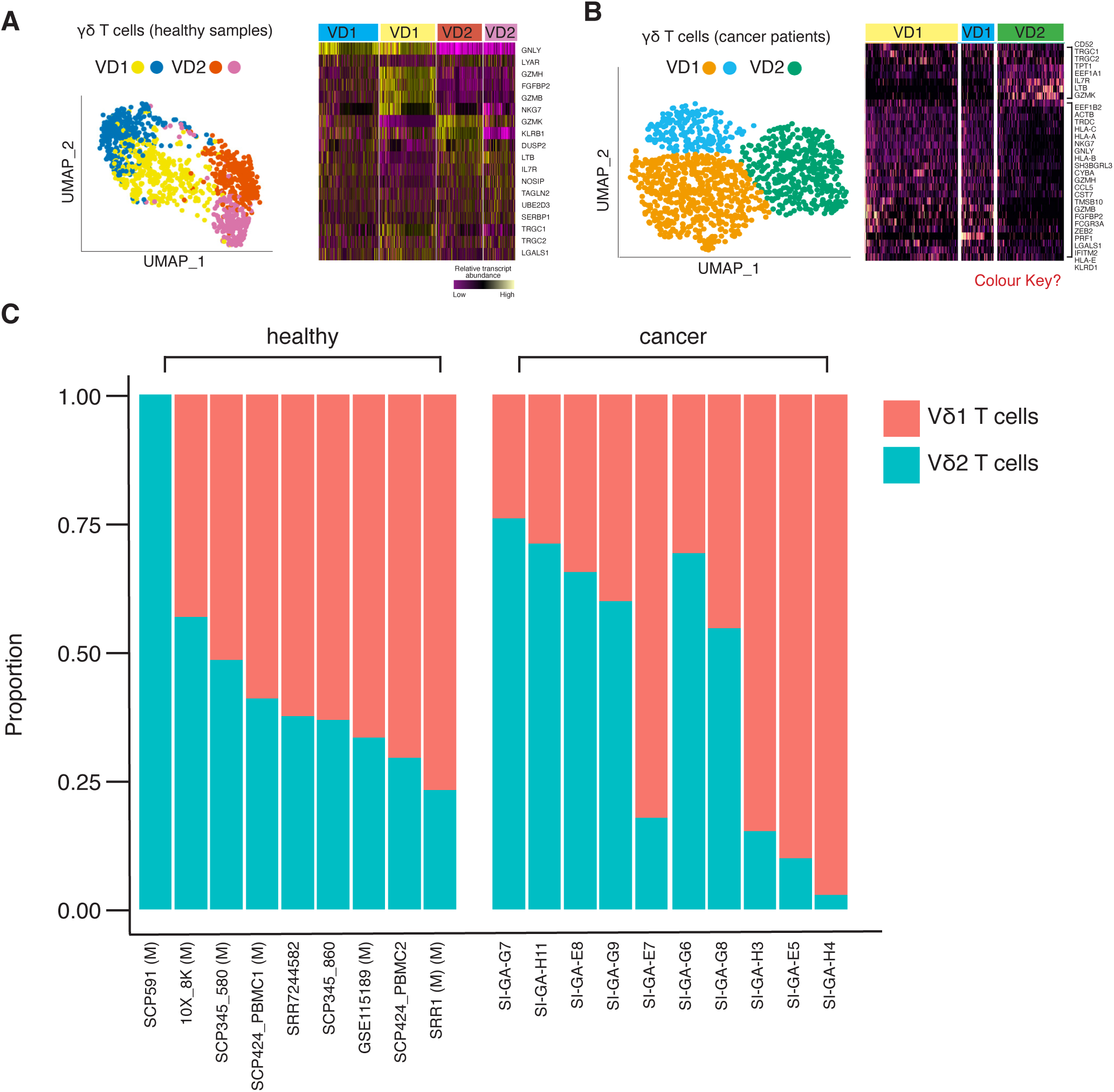
Circulating Vδ 1 and Vδ 2 subsets show phenotypic changes in metastatic breast cancer. **A-B.** UMAP plots of the re-clustered γδ T cells from PBMCs of healthy and metastatic breast cancer cohorts, coloured by the relative transcript abundance of the top marker genes. The heatmaps show top differentially abundant transcripts in γδ T subsets in healthy (**A**) and cancer (**C**) PBMC samples. Heatmaps are coloured by the relative abundance of marker-gene transcripts (row-scaled). **C.** The plot shows the relative proportion of Vδ 1 and Vδ 2 T cell subtypes in healthy and metastatic cancer patients (including low metastatic burden cohort).

## References

1. Fares, J., et al., Molecular principles of metastasis: a hallmark of cancer revisited. Signal Transduct Target Ther, 2020. 5(1): p. 28.

2. Riggio, A.I., K.E. Varley, and A.L. Welm, The lingering mysteries of metastatic recurrence in breast cancer. Br J Cancer, 2021. 124(1): p. 13–26.

3. Waks, A.G., et al., The Immune Microenvironment in Hormone Receptor-Positive Breast Cancer Before and After Preoperative Chemotherapy. Clin Cancer Res, 2019. 25(15): p. 4644–4655.

4. Keller, L., et al., Clinical relevance of blood-based ctDNA analysis: mutation detection and beyond. Br J Cancer, 2021. 124(2): p. 345–358.

5. Pang, S., et al., Circulating tumour cells at baseline and late phase of treatment provide prognostic value in breast cancer. Sci Rep, 2021. 11(1): p. 13441.

6. Gracie, L., et al., Circulating tumour DNA (ctDNA) in metastatic melanoma, a systematic review and meta-analysis. Eur J Cancer, 2021. 158: p. 191–207.

7. Gonzalez, H., I. Robles, and Z. Werb, Innate and acquired immune surveillance in the postdissemination phase of metastasis. FEBS J, 2018. 285(4): p. 654–664.

8. Gonzalez, H., C. Hagerling, and Z. Werb, Roles of the immune system in cancer: from tumor initiation to metastatic progression. Genes Dev, 2018. 32(19-20): p. 1267–1284.

9. Twine, N.C., et al., Disease-associated expression profiles in peripheral blood mononuclear cells from patients with advanced renal cell carcinoma. Cancer Res, 2003. 63(18): p. 6069–75.

10. Showe, M.K., et al., Gene expression profiles in peripheral blood mononuclear cells can distinguish patients with non-small cell lung cancer from patients with nonmalignant lung disease. Cancer Res, 2009. 69(24): p. 9202–10.

11. Showe, M.K., A.V. Kossenkov, and L.C. Showe, The peripheral immune response and lung cancer prognosis. Oncoimmunology, 2012. 1(8): p. 1414–1416.

12. Patarat, R., et al., The expression of FLNA and CLU in PBMCs as a novel screening marker for hepatocellular carcinoma. Sci Rep, 2021. 11(1): p. 14838.

13. Foulds, G.A., et al., Immune-Phenotyping and Transcriptomic Profiling of Peripheral Blood Mononuclear Cells From Patients With Breast Cancer: Identification of a 3 Gene Signature Which Predicts Relapse of Triple Negative Breast Cancer. Front Immunol, 2018. 9: p. 2028.

14. Ming, W., et al., Two Distinct Subtypes Revealed in Blood Transcriptome of Breast Cancer Patients With an Unsupervised Analysis. Front Oncol, 2019. 9: p. 985.

15. Lun, Y.Z., J. Sun, and Z.G. Yu, [Identification of Differentially Expressed Gene Core Genes in Early T-Cell Precursor Acute Lymphoblastic Leukemia and Its Regulatory Network Analysis]. Zhongguo Shi Yan Xue Ye Xue Za Zhi, 2019. 27(3): p. 673–684.

16. Hao, Y., et al., Integrated analysis of multimodal single-cell data. Cell, 2021. 184(13): p. 3573–3587 e29.

17. Freytag, S., et al., Comparison of clustering tools in R for medium-sized 10x Genomics single-cell RNA-sequencing data. F1000Res, 2018. 7: p. 1297.

18. Ding, J., et al., Systematic comparison of single-cell and single-nucleus RNA-sequencing methods. Nat Biotechnol, 2020. 38(6): p. 737–746.

19. Karagiannis, T.T., et al., Single cell transcriptomics reveals opioid usage evokes widespread suppression of antiviral gene program. Nat Commun, 2020. 11(1): p. 2611.

20. Cai, Y., et al., Single-cell transcriptomics of blood reveals a natural killer cell subset depletion in tuberculosis. EBioMedicine, 2020. 53: p. 102686.

21. Gold, M.C., et al., Human mucosal associated invariant T cells detect bacterially infected cells. PLoS Biol, 2010. 8(6): p. e1000407.

22. Kurioka, A., et al., MAIT cells are licensed through granzyme exchange to kill bacterially sensitized targets. Mucosal Immunol, 2015. 8(2): p. 429–40.

23. Gherardin, N.A., et al., Enumeration, functional responses and cytotoxic capacity of MAIT cells in newly diagnosed and relapsed multiple myeloma. Sci Rep, 2018. 8(1): p. 4159.

24. Pal, B., et al., A single-cell RNA expression atlas of normal, preneoplastic and tumorigenic states in the human breast. EMBO J, 2021. 40(11): p. e107333.

25. Bouwman, A.C., et al., Intracellular and Extracellular Roles of Granzyme K. Front Immunol, 2021. 12: p. 677707.

26. Tibbs, E. and X. Cao, Emerging Canonical and Non-Canonical Roles of Granzyme B in Health and Disease. Cancers (Basel), 2022. 14(6).

27. Tewary, P., et al., Granulysin activates antigen-presenting cells through TLR4 and acts as an immune alarmin. Blood, 2010. 116(18): p. 3465–74.

28. Hermani, A., et al., Calcium-binding proteins S100A8 and S100A9 as novel diagnostic markers in human prostate cancer. Clin Cancer Res, 2005. 11(14): p. 5146–52.

29. Allgower, C., et al., Friend or Foe: S100 Proteins in Cancer. Cancers (Basel), 2020. 12(8).

30. Ntanasis-Stathopoulos, I., D. Fotiou, and E. Terpos, CCL3 Signaling in the Tumor Microenvironment. Adv Exp Med Biol, 2020. 1231: p. 13–21.

31. Baker, K.J., A. Houston, and E. Brint, IL-1 Family Members in Cancer; Two Sides to Every Story. Front Immunol, 2019. 10: p. 1197.

32. Guo, H., et al., SCARB2/LIMP-2 Regulates IFN Production of Plasmacytoid Dendritic Cells by Mediating Endosomal Translocation of TLR9 and Nuclear Translocation of IRF7. J Immunol, 2015. 194(10): p. 4737–49.

33. Morrell, C.N., et al., Emerging roles for platelets as immune and inflammatory cells. Blood, 2014. 123(18): p. 2759–67.

34. Salemme, V., et al., The Crosstalk Between Tumor Cells and the Immune Microenvironment in Breast Cancer: Implications for Immunotherapy. Front Oncol, 2021. 11: p. 610303.

35. Wu, B., et al., Cross-talk between cancer stem cells and immune cells: potential therapeutic targets in the tumor immune microenvironment. Mol Cancer, 2023. 22(1): p. 38.

36. Dias, A.S., et al., Metabolic crosstalk in the breast cancer microenvironment. Eur J Cancer, 2019. 121: p. 154–171.

37. Jin, S., et al., Inference and analysis of cell-cell communication using CellChat. Nat Commun, 2021. 12(1): p. 1088.

38. Stein, E.V., et al., Secreted Thrombospondin-1 Regulates Macrophage Interleukin-1beta Production and Activation through CD47. Sci Rep, 2016. 6: p. 19684.

39. Kumanogoh, A., et al., Nonredundant roles of Sema4A in the immune system: defective T cell priming and Th1/Th2 regulation in Sema4A-deficient mice. Immunity, 2005. 22(3): p. 305–16.

40. Godfrey, D.I., et al., The biology and functional importance of MAIT cells. Nat Immunol, 2019. 20(9): p. 1110–1128.

41. Toubal, A., et al., Mucosal-associated invariant T cells and disease. Nat Rev Immunol, 2019. 19(10): p. 643–657.

42. Barrow, A.D., C.J. Martin, and M. Colonna, The Natural Cytotoxicity Receptors in Health and Disease. Front Immunol, 2019. 10: p. 909.

43. Fujii, S., et al., Arl4c expression in colorectal and lung cancers promotes tumorigenesis and may represent a novel therapeutic target. Oncogene, 2015. 34(37): p. 4834–44.

44. Su, L., R.J. Hershberger, and I.L. Weissman, LYAR, a novel nucleolar protein with zinc finger DNA-binding motifs, is involved in cell growth regulation. Genes Dev, 1993. 7(5): p. 735–48.

45. Wu, Y., et al., LYAR promotes colorectal cancer cell mobility by activating galectin-1 expression. Oncotarget, 2015. 6(32): p. 32890–901.

46. Lang, R. and F.A.M. Raffi, Dual-Specificity Phosphatases in Immunity and Infection: An Update. Int J Mol Sci, 2019. 20(11).

47. Upadhyay, V. and Y.X. Fu, Lymphotoxin signalling in immune homeostasis and the control of microorganisms. Nat Rev Immunol, 2013. 13(4): p. 270–9.

48. Fei, F., et al., Role of metastasis-induced protein S100A4 in human non-tumor pathophysiologies. Cell Biosci, 2017. 7: p. 64.

49. Pizzolato, G., et al., Single-cell RNA sequencing unveils the shared and the distinct cytotoxic hallmarks of human TCRVdelta1 and TCRVdelta2 gammadelta T lymphocytes. Proc Natl Acad Sci U S A, 2019. 116(24): p. 11906–11915.

50. Ng, S.S., et al., The NK cell granule protein NKG7 regulates cytotoxic granule exocytosis and inflammation. Nat Immunol, 2020. 21(10): p. 1205–1218.

51. Yim, C.Y., et al., G0S2 Suppresses Oncogenic Transformation by Repressing a MYC-Regulated Transcriptional Program. Cancer Res, 2016. 76(5): p. 1204–13.

52. Cho, E., et al., G0/G1 Switch 2 Induces Cell Survival and Metastasis through Integrin-Mediated Signal Transduction in Human Invasive Breast Cancer Cells. Biomol Ther (Seoul), 2019: p. 591–602.

53. Okabe, M., et al., G0S2 regulates innate immunity in Kawasaki disease via lncRNA HSD11B1-AS1. Pediatr Res, 2022.

54. Fujikawa, M., et al., Population of ATP synthase molecules in mitochondria is limited by available 6.8-kDa proteolipid protein (MLQ). Genes Cells, 2014. 19(2): p. 153–60.

55. Klein, K., et al., Role of Mitochondria in Cancer Immune Evasion and Potential Therapeutic Approaches. Front Immunol, 2020. 11: p. 573326.

56. Becuwe, P., et al., Manganese superoxide dismutase in breast cancer: from molecular mechanisms of gene regulation to biological and clinical significance. Free Radic Biol Med, 2014. 77: p. 139–51.

57. Chen, Y., et al., CD36, a signaling receptor and fatty acid transporter that regulates immune cell metabolism and fate. J Exp Med, 2022. 219(6).

58. Kang, S.W., Superoxide dismutase 2 gene and cancer risk: evidence from an updated meta-analysis. Int J Clin Exp Med, 2015. 8(9): p. 14647–55.

59. Billard, M.J., et al., G Protein Coupled Receptor Kinase 3 Regulates Breast Cancer Migration, Invasion, and Metastasis. PLoS One, 2016. 11(4): p. e0152856.

60. Li, W., et al., GRK3 is essential for metastatic cells and promotes prostate tumor progression. Proc Natl Acad Sci U S A, 2014. 111(4): p. 1521–6.

61. Weinelt, N. and S.J.L. van Wijk, Ubiquitin-dependent and -independent functions of OTULIN in cell fate control and beyond. Cell Death Differ, 2021. 28(2): p. 493–504.

62. Heger, K., et al., OTULIN limits cell death and inflammation by deubiquitinating LUBAC. Nature, 2018. 559(7712): p. 120–124.

63. Moradpoor, R., et al., Identification and Validation of Stage-Associated PBMC Biomarkers in Breast Cancer Using MS-Based Proteomics. Front Oncol, 2020. 10: p. 1101.

64. Zimmerman, A.W., et al., Long-term engagement of CD6 and ALCAM is essential for T-cell proliferation induced by dendritic cells. Blood, 2006. 107(8): p. 3212–20.

65. Hinks, T.S.C. and X.W. Zhang, MAIT Cell Activation and Functions. Front Immunol, 2020. 11: p. 1014.

66. Jiang, X., N.K. Bjorkstrom, and E. Melum, Intact CD100-CD72 Interaction Necessary for TCR-Induced T Cell Proliferation. Front Immunol, 2017. 8: p. 765.

67. Couty, J.P., et al., PECAM-1 engagement counteracts ICAM-1-induced signaling in brain vascular endothelial cells. J Neurochem, 2007. 103(2): p. 793–801.

68. Rajasekaran, K., et al., Signaling in Effector Lymphocytes: Insights toward Safer Immunotherapy. Front Immunol, 2016. 7: p. 176.

69. Orcajo-Rincon, J., et al., Review of imaging techniques for evaluating morphological and functional responses to the treatment of bone metastases in prostate and breast cancer. Clin Transl Oncol, 2022. 24(7): p. 1290–1310.

70. Kwee, T.C. and R.M. Kwee, Combined FDG-PET/CT for the detection of unknown primary tumors: systematic review and meta-analysis. Eur Radiol, 2009. 19(3): p. 731–44.

71. Lother, D., et al., Imaging in metastatic breast cancer, CT, PET/CT, MRI, WB-DWI, CCA: review and new perspectives. Cancer Imaging, 2023. 23(1): p. 53.

72. Zheng, G.X., et al., Massively parallel digital transcriptional profiling of single cells. Nat Commun, 2017. 8: p. 14049.

73. Amezquita, R.A., et al., Orchestrating single-cell analysis with Bioconductor. Nat Methods, 2020. 17(2): p. 137–145.

74. Lun, A.T.L., et al., EmptyDrops: distinguishing cells from empty droplets in droplet-based single-cell RNA sequencing data. Genome Biol, 2019. 20(1): p. 63.

75. McCarthy, D.J., et al., Scater: pre-processing, quality control, normalization and visualization of single-cell RNA-seq data in R. Bioinformatics, 2017. 33(8): p. 1179–1186.

76. Mangiola, S., M.A. Doyle, and A.T. Papenfuss, Interfacing Seurat with the R tidy universe. Bioinformatics, 2021.

77. McGinnis, C.S., L.M. Murrow, and Z.J. Gartner, DoubletFinder: Doublet Detection in Single-Cell RNA Sequencing Data Using Artificial Nearest Neighbors. Cell Syst, 2019. 8(4): p. 329–337 e4.

78. Butler, A., et al., Integrating single-cell transcriptomic data across different conditions, technologies, and species. Nat Biotechnol, 2018. 36(5): p. 411–420.

79. Stuart, T., et al., Comprehensive Integration of Single-Cell Data. Cell, 2019. 177(7): p. 1888–1902 e21.

80. Xu, C. and Z. Su, Identification of cell types from single-cell transcriptomes using a novel clustering method. Bioinformatics, 2015. 31(12): p. 1974–80.

81. Cetintas, S., et al., Prediction of breast cancer metastasis risk using circulating tumor markers: A follow-up study. Bosn J Basic Med Sci, 2019. 19(2): p. 172–179.

82. Mangiola, S., et al., sccomp: Robust differential composition and variability analysis for single-cell data. Proc Natl Acad Sci U S A, 2023. 120(33): p. e2203828120.

83. Mangiola, S., et al., tidybulk: an R tidy framework for modular transcriptomic data analysis. Genome Biol, 2021. 22(1): p. 42.

84. Zhou, X., H. Lindsay, and M.D. Robinson, Robustly detecting differential expression in RNA sequencing data using observation weights. Nucleic Acids Res, 2014. 42(11): p. e91.

85. McCarthy, D.J. and G.K. Smyth, Testing significance relative to a fold-change threshold is a TREAT. Bioinformatics, 2009. 25(6): p. 765–71.

86. Mangiola, S., et al., Probabilistic outlier identification for RNA sequencing generalized linear models. NAR Genom Bioinform, 2021. 3(1): p. lqab005.

87. Robinson, M.D. and A. Oshlack, A scaling normalization method for differential expression analysis of RNA-seq data. Genome Biol, 2010. 11(3): p. R25.

88. Bunis, D.G., et al., dittoSeq: Universal User-Friendly Single-Cell and Bulk RNA Sequencing Visualization Toolkit. Bioinformatics, 2020.

89. Wickham, H., ggplot2: Elegant Graphics for Data Analysis. 2016: Springer International Publishing.

90. S, M. and P. A., T. tidyHeatmap: an R package for modular heatmap production based on tidy principles. Journal of Open Source Software 2020. 5(52).

91. Wickham, H., et al., Welcome to the Tidyverse. Journal of Open Source Software, 2019.

